# Genetic association meta-analysis is susceptible to confounding by between-study cryptic relatedness

**DOI:** 10.1101/2025.05.10.653279

**Authors:** Tiffany Tu, Alejandro Ochoa

## Abstract

Meta-analysis of Genome-Wide Association Studies (GWAS) has important advantages, but it assumes that studies are independent, which does not hold when there is relatedness between studies. As a motivating example, recent work suggested applying sex-stratified meta-analysis to correct for participation bias, without considering that men and women from the same population will be highly related. Our theory demonstrates how cryptic relatedness results in correlated test statistics between studies, inflating meta-analysis. We characterize the effects of different between-study relatedness scenarios, particularly population structure and recent family relatedness, on meta-analysis type I error control and power. We simulated data with (1) no family relatedness between subpopulations, (2) family relatedness within subpopulations, (3) family relatedness across subpopulations, and (4) single population with family relatedness. We run joint and meta-analyses on simulations using both binary and quantitative traits. In scenarios with family relatedness, sex-stratified meta-analysis exhibits severe inflation and lower AUC compared to joint and subpopulation meta-analyses. Remarkably, genomic control succeeds in correcting inflation in these cases, but does not alter calibrated power. Analysis of real datasets confirms severe inflation for sex-stratified meta-analysis in family studies, but a negligible effect for population studies with up to 10,000 individuals. Our theoretical framework demonstrates that the inflation factor increases as the sample size increases. We recommend against meta-analyzing studies that share the same populations, which increases the risk of inflation due to cryptic relatedness between studies.

## 1 Introduction

Meta-analysis of Genome-Wide Association Studies (GWAS) combines summary statistics from multiple studies, which routinely increases power compared to a single GWAS, permits analysis without sharing individual-level data, and better accounts for potential heterogeneity between different populations, among other advantages [1, 2] (Figure 1A). Population structure due to ancestry differences and cryptic relatedness, which can be understood as ancient and recent (family) relatedness that are unknown to researchers, respectively (Figure 1B), are important confounders in GWAS when they are not adequately modeled [3, 4]. Modern GWAS successfully model population structure with PCA covariates [5] and address both population structure and cryptic relatedness with linear and logistic mixed-effects models [6–12]. However, some meta-analysis designs can result in non-negligible cryptic relatedness between studies, which is an additional confounding factor that cannot be adequately addressed by modeling relatedness solely within studies. Cryptic relatedness between studies violates the independence assumption that meta-analysis makes, as we show here, leading to correlated errors between test statistics of different studies, resulting in increased type I error rates in the meta-analysis. Thus, confounding can occur in the meta-analysis even when the input studies are not individually confounded.

**Fig. 1:**
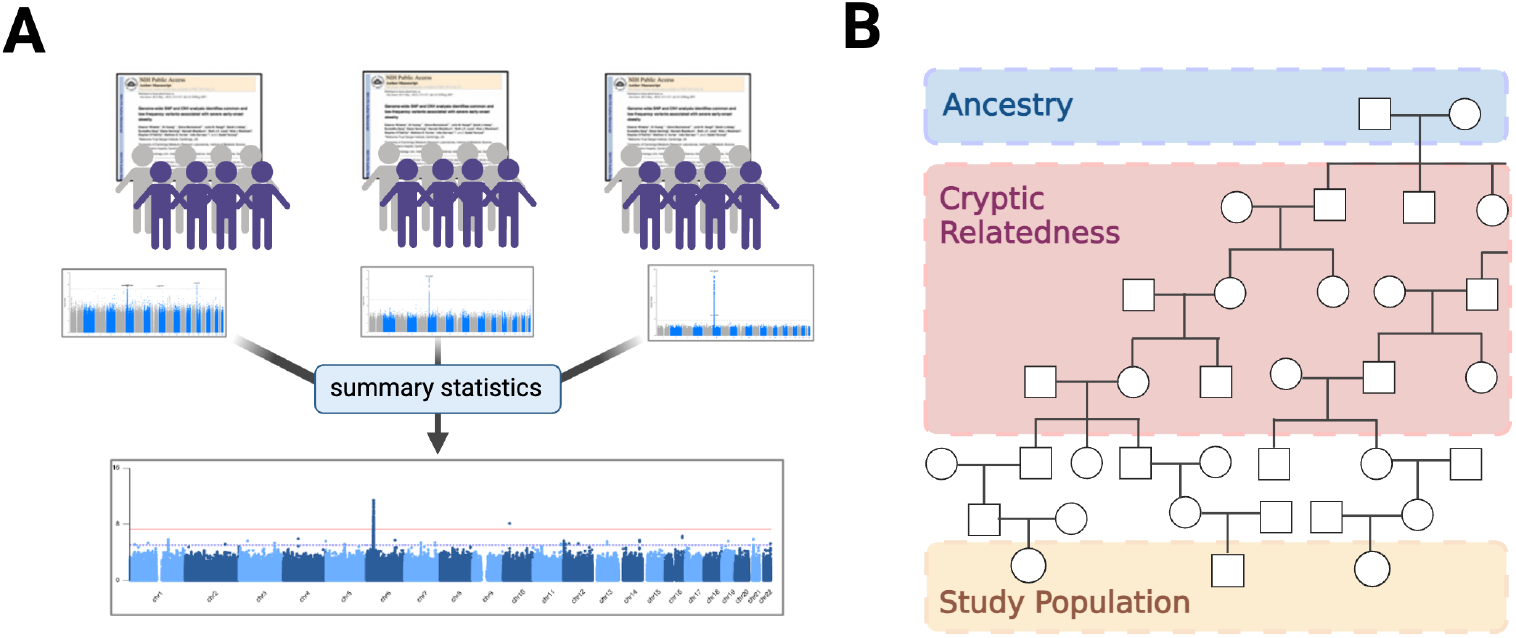
Cryptic relatedness can cause inflation in GWAS meta analysis. **A**. Illustration of a standard GWAS meta-analysis pipeline, which combines summary statistics from multiple studies while assuming data independence. Population structure due to ancestry differences and cryptic relatedness are both common causes of dependence between individuals in GWAS. **B**. Different degrees of relatedness visualized through a pedigree. Cryptic relatedness is a form of family relatedness that is recent but unknown to the researchers. Family structure is high dimensional, and must be modeled with linear mixed-effects models in GWAS [12]. In contrast, ancestry is a more ancient form of relatedness that is broadly shared across the population, so it can be modeled with low dimensional models such as PCA. (Created with BioRender.com)

The importance of sample independence is well known to statistical geneticists, and it is recommended to adjust for population stratification and to identify sample overlap between studies before carrying out meta-analysis [1]. Despite these guidelines, there is still limited attention on addressing cryptic relatedness between studies in meta-analyses. For example, a recent study recommended performing sex-stratified meta-analysis to correct for sex-differential participation bias, without considering the potential violation of independence between the male and female studies [13]. Meta-analysis does not allow direct modeling of relatedness between studies, but there are potential alternatives. One is to adjust output statistics using Genomic Control (GC), which calibrates the median statistic to its expected value [3, 9, 14–17]. However, while GC was originally proposed to correct for population stratification, it is often unsuccessful [17–19], and it has been studied even less for correcting for cryptic relatedness. Another is joint GWAS of the substudies (or mega-analysis), which effectively handles population structure and cryptic relatedness, but it requires access to individual data, which can be a major hindrance. Federated learning privacy approaches are recent proposals addressing this problem, which perform joint analyses of data across institutes without sharing individual-level data, instead performing calculations using encrypted genotypes to both detect genetic relatedness across cohorts [20–22] and perform association tests [23–25]. A recent study compared joint (or mega) GWAS to meta-analysis in a multi-ancestry setting using data from the Hyperglycemia and Adverse Pregnancy Outcome (HAPO) study, and found that joint GWAS has more variability in genomic inflation but improved power and identified more significant genetic associations [26]. Simulations and real data from UK Biobank and All of Us Research Programs demonstrate that joint (or pooled) analysis exhibits better statistical power while effectively adjusting for population stratification [27].

In this work, we demonstrate that cryptic relatedness between studies can cause confounding in meta-analysis. We simulate data with combinations of population and family structure, and identify family structure as the only severe source of confounding. Remarkably, we find that GC can successfully correct inflated meta-analysis results, although loss of power remains unchanged. Analysis of real human datasets confirms our findings, with strong confounding observed in family studies, although the effect is negligible in population studies. Nevertheless, we expect cryptic relatedness to become a more severe confounder of meta-analyses at ever larger sample sizes.

## 2 Methods

### 2.1 Genetic association method

Joint and stratified GWAS are performed using SAIGE (Scalable and Accurate Implementation of GEn-eralized mixed model), which controls for unbalanced case-control ratio and sample relatedness at efficient runtime and scalability [10]. The approach for quantitative traits is a linear model (shown below), while for binary traits a logistic mixed model is used instead (not shown). Each of our GWAS incorporates as covariates sex, age (for real datasets), and subpopulation label (for simulated datasets), if they are not constant, along with the top 10 principal components (PCs) of the genotype matrix. We use METAL [28] to perform all of our fixed-effects GWAS meta-analyses.

The linear mixed-effects model (LMM) used at SNP *i* for a given study *j* with *n*_*j*_ individuals is given by

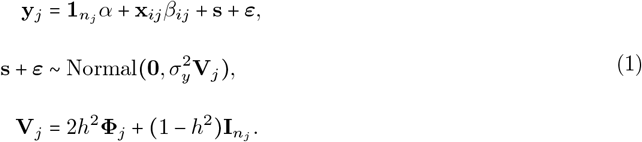

where **y**_*j*_ is a length-*n*_*j*_ vector of individual trait values, 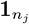 is a length-*n*_*j*_ vector of ones, *α* is a scalar of intercept coefficients, **x**_*ij*_ is the length-*n*_*j*_ vector of genotypes at SNP *i, β*_*ij*_ is the genetic effect coefficient, **s** is a length *n*_*j*_ structured residual vector, ***ε*** is a length-*n*_*j*_ vector of residual errors, 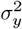 is a trait-specific variance factor, **V**_*j*_ is the *n*_*j*_ × *n*_*j*_ scale-free covariance matrix of the trait, *h*^2^ is the heritability, **Φ**_*j*_ is the *n*_*j*_ × *n*_*j*_ kinship matrix and 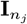 is the *n*_*j*_ × *n*_*j*_ identity matrix. The genetic effect coefficient 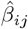 and its standard error 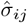 are estimated from this model as shown in Appendix A.

### 2.2 Inflation factor and the Genomic Control method

A common metric to assess confounding due to population stratification in GWAS, or statistic miscalibration more broadly, is the inflation factor *λ*, defined as the median test statistic across SNPs divided by the theoretical median under the expected null *χ*^2^ distribution [3, 9, 14–17]. Calibrated statistics have *λ* ≈ 1, while *λ* >1.05 is considered inflated [9]. The Genomic Control (GC) method consists of dividing test statistics by the inflation factor, thus ensuring that the inflation factor is exactly one after the correction [3], although this does not guarantee calibration at other quantiles. Despite this potential limitation, we consider GC as a possible solution to meta-analysis inflation caused by relatedness between studies.

### 2.3 Genotype and trait simulations

Genotypes with population and family structure are simulated using the R packages `bnpsd` and `simfam`, respectively, following previous work [12, 29]. Briefly, an ancestral population *T* has allele frequencies for 500,000 SNPs drawn from a uniform distribution, then subpopulations *S*_*i*_ for *i* ∈ {1, 2, 3} have allele frequencies drawn from the Balding-Nichols model [30], from which genotypes are drawn binomially for 1,000 individuals per subpopulation (3,000 in total). We generate correlated *S*_2_ and *S*_3_ by drawing them from a common ancestor population, denoted *S*_23_, while *S*_1_ is simulated independently. Subpopulations *S*_1_ and *S*_23_ both have an inbreeding coefficient from *T* of 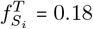. Subpopulations *S*_2_ and *S*_3_ have 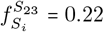 from their common ancestor, resulting in a total inbreeding coefficient from *T* of 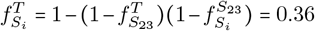 (formula from [29]). The average inbreeding coefficient across the combined population is *F*_ST_ = 0.3. Sex is drawn randomly for each individual.

When there is family structure, pedigrees are constructed by pairing individuals of opposite sexes randomly biased by proximity in their one-dimensional coordinates (initially contiguous by subpopulation), with variable numbers of children constrained to a total desired population size, which is fixed throughout the generations. Founders have genotypes drawn from the same population structure described above. Child genotypes are drawn from their parents genotypes following Mendelian inheritance, independently per locus. To characterize the effect of between-study relatedness scenarios, we considered four simulations: (1) no family relatedness across 3 subpopulations; or otherwise 30 generations of family relatedness with these configurations: (2) within subpopulations, (3) across subpopulations, and (4) in a single population (Figure 2). Since simulation 4 has a single population (with 3000 individuals in total, matching the rest), for testing subpopulation-stratified meta-analysis, it is partitioned into three “subpopulations” of 1000 individuals each grouped by their one-dimensional coordinates.

**Fig. 2:**
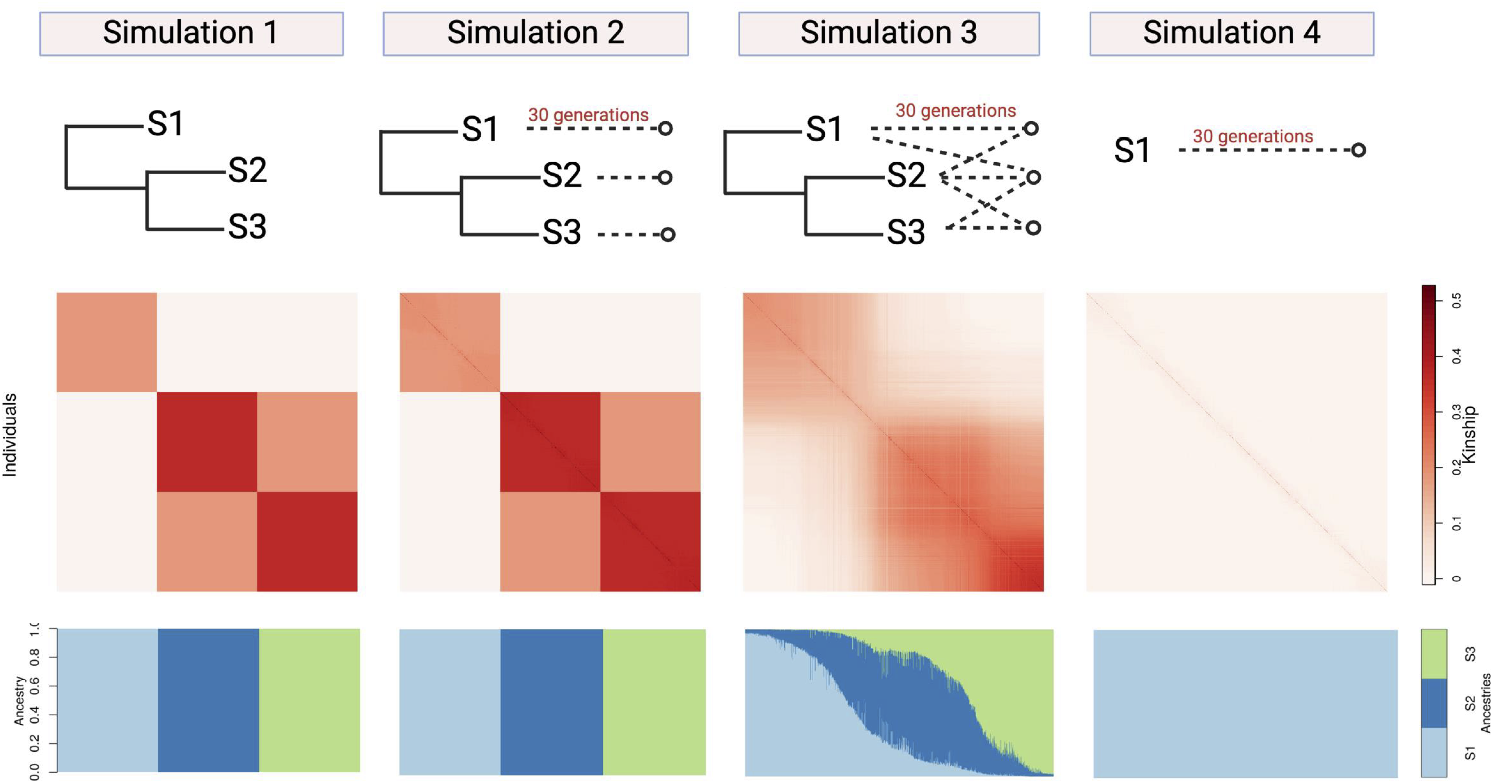
Overview of the population and family structure of simulation scenarios. **Top row:** Phylogenetic trees summarize the key features of each relatedness structure. Population and family structure are represented by solid and dashed lines, respectively. **Middle row:** Kinship matrices show the covariance structure between each pair of individuals (along both x and y axes, using the same order of the underlying one-dimensional space of the simulations), where color represents their total kinship coefficient (reflecting both population and family relatedness). Diagonal plots inbreeding coefficients instead of self kinship since inbreeding is in the same scale as the rest of kinship values. Note family structure results in more high-kinship pairs that appear near the diagonal of simulations 2-4. **Bottom row:** Admixture proportions better summarize the population structure in this data, ignoring family structure. Each individual (x axis) has a stacked barplot of ancestry proportions (colors) that sums to one. Only simulation 3 has admixture (individuals with more than one ancestry).

Quantitative traits are simulated using R package `simtrait`, follow a fixed effect size (FES) model for the coefficients with *h*^2^ = 0.8 and 100 causal loci, as before [12]. Binary traits are generated by thresholding the previous quantitative traits at their median. Traits are resimulated for a replicate if SAIGE does not converge, which occurs if case-control ratios in a given subpopulation are extremely imbalanced.

### 2.4 Evaluation metrics

Our simulations are evaluated using (1) the inflation factor *λ* defined earlier, (2) AUC_PR_ (precision-recall area under the curve), and (3) SRMSD_*p*_ (p-value signed root mean square deviation), following earlier work [12]. Briefly, we use the R package `PRROC` [31] to compute AUC_PR_, where given the total number of true positives (TP), false positives (FP), and false negatives (FN) at some p-value threshold *t*,

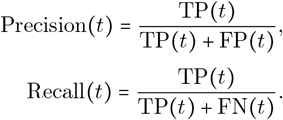

The resulting area under the curve, AUC_PR_, measures causal locus classification accuracy and reflects calibrated statistical power while robust to miscalibrated models [12].

SRMSD_*p*_ measures the difference between the observed null p-value quantiles and the expected uniform quantiles:

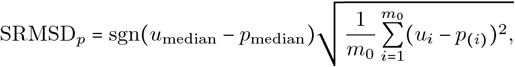

where *m*_0_ is the number of null (non-causal) loci, *p*_(*i*)_ is the *i*th order null p-value, *u*_*i*_ = (*i* − 0.5)/*m*_0_ is its expectation, *p*_median_ is the median observed null p-value, 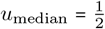 is its expectation, and sgn is the sign function. Analogous to the 0.95 < *λ* < 1.05 rule of thumb, ∣SRMSD_*p*_∣ < 0.01 is considered calibrated [12].

SRMSD_*p*_ is a stricter test of calibration than the inflation factor, since the ideal SRMSD_*p*_ =0 requires null p-value uniformity at all quantiles, whereas the ideal *λ* = 1 merely requires the median to match its null expectation. However, since both AUC_PR_ and SRMSD_*p*_ require knowing truly causal loci, they cannot be applied to real data. In contrast, inflation factor can be computed directly from the observed summary statistics in real datasets.

### 2.5 Real datasets

#### T2D-GENES SAMAFS Consortium

We analyzed 914 individuals (380 males and 534 females) for 21 traits from the San Antonio Mexican American Family Studies (SAMAFS) Project 2, using exome chip data to maximize the number of individuals from 20 large Mexican American pedigrees (dbGaP accession phs000847.v2.p1). Starting from 79,272 SNPs, we applied filters of Hardy-Weinberg equilibrium (HWE) *p* < 1e-10 and a minor allele frequency (MAF) < 0.01, resulting in 36,293 SNPs. A total of 20 quantitative traits and one binary trait (Type 2 Diabetes) were analyzed (Table 1). Quantitative traits were log-transformed where appropriate to improve normality.

**Table 1:**
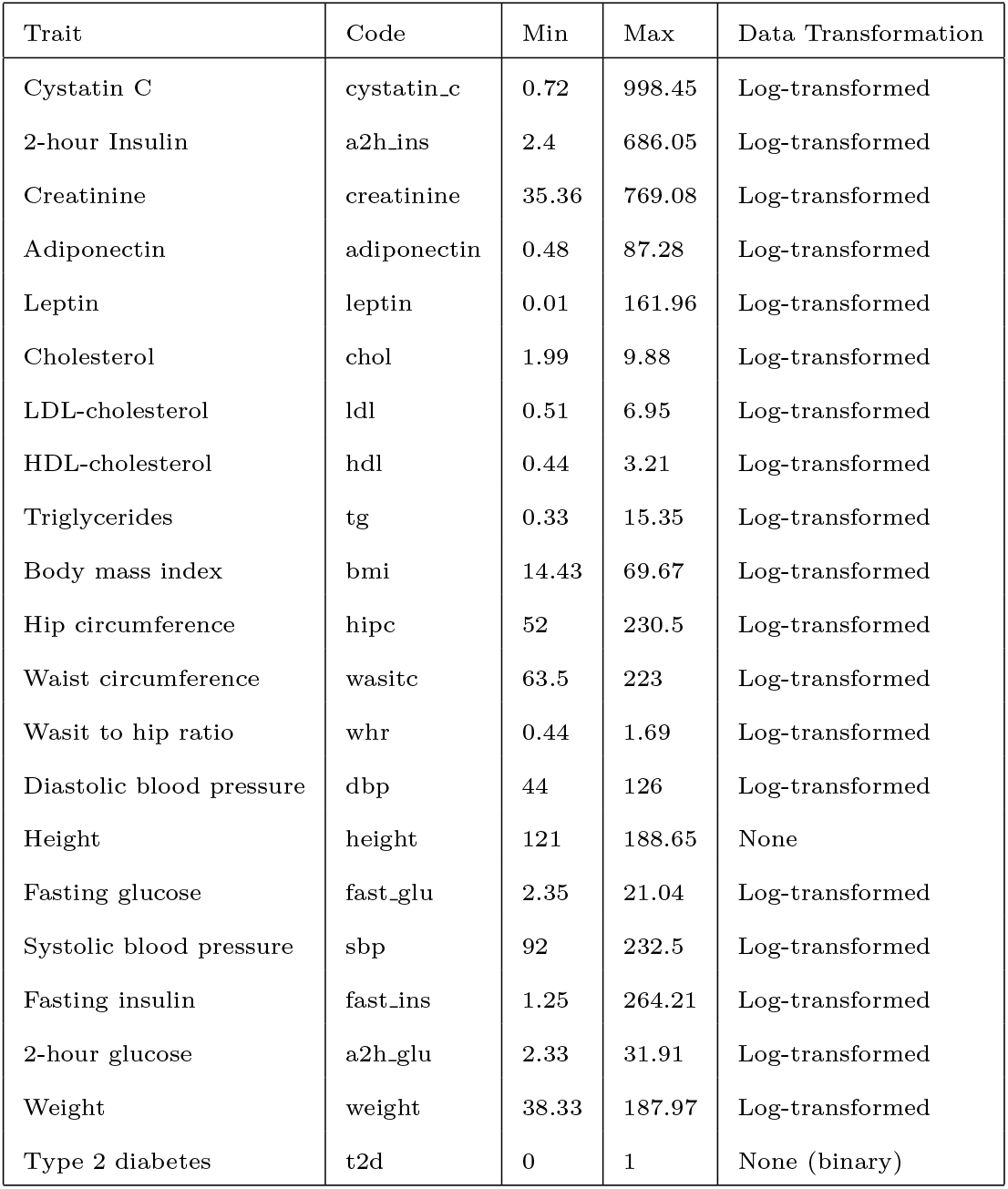
Traits included in SAMAFS analysis, including data transformation applied.

| Trait | Code | Min | Max | Data Transformation |
| --- | --- | --- | --- | --- |
| Cystatin C | cystatin_c | 0.72 | 998.45 | Log-transformed |
| 2-hour Insulin | a2h_ins | 2.4 | 686.05 | Log-transformed |
| Creatinine | creatinine | 35.36 | 769.08 | Log-transformed |
| Adiponectin | adiponectin | 0.48 | 87.28 | Log-transformed |
| Leptin | leptin | 0.01 | 161.96 | Log-transformed |
| Cholesterol | chol | 1.99 | 9.88 | Log-transformed |
| LDL-cholesterol | ldl | 0.51 | 6.95 | Log-transformed |
| HDL-cholesterol | hdl | 0.44 | 3.21 | Log-transformed |
| Triglycerides | tg | 0.33 | 15.35 | Log-transformed |
| Body mass index | bmi | 14.43 | 69.67 | Log-transformed |
| Hip circumference | hipc | 52 | 230.5 | Log-transformed |
| Waist circumference | wasitc | 63.5 | 223 | Log-transformed |
| Wasit to hip ratio | whr | 0.44 | 1.69 | Log-transformed |
| Diastolic blood pressure | dbp | 44 | 126 | Log-transformed |
| Height | height | 121 | 188.65 | None |
| Fasting glucose | fast_glu | 2.35 | 21.04 | Log-transformed |
| Systolic blood pressure | sbp | 92 | 232.5 | Log-transformed |
| Fasting insulin | fast_ins | 1.25 | 264.21 | Log-transformed |
| 2-hour glucose | a2h_glu | 2.33 | 31.91 | Log-transformed |
| Weight | weight | 38.33 | 187.97 | Log-transformed |
| Type 2 diabetes | t2d | 0 | 1 | None (binary) |

#### HCHS/SOL

We analyzed 11,721 individuals (6831 males and 4890 females) from The Hispanic Community Healthy Study/Study of Latinos (HCHS/SOL) (dbGaP accession phs000810.v2.p2) using array data from substudy `SOL_phaseIa_genotyping.` We applied HWE *p* < 1e-10 and MAF < 0.01 filters, resulting in 1,651,164 SNPs. A total of 33 quantitative traits were analyzed and transformed as needed (Table 2).

**Table 2:** Traits included in HCHS/SOL analysis, including data transformation applied.

| Trait | Code | Units | Min | Max | Data Transformation |
| --- | --- | --- | --- | --- | --- |
| Height | HEIGHT | cm | 122 | 199 | None |
| Body mass index | BMI | kg/m2 | 14.28 | 70.35 | Log-transformed |
| Waist to hip ratio | WASIT_HIP | NA | 0.52 | 1.31 | Log-transformed |
| Insulin fasting | INSULIN_FAST | calibrated to mU/L | 0.33 | 726 | Log-transformed |
| Glucose fasting | LABA70 | mg/dL | 52 | 642 | Log-transformed |
| Glucose post Oral Glucose Tolerance Test (OGTT) | LABA71 | mg/dL | 29 | 389 | Log-transformed |
| Cystatin c. | LABA101 | mg/L | 0.34 | 10.1 | Log-transformed |
| High-sensitivity c-reactive protein | LABA91 | mg/L | 0.11 | 316.9 | Log-transformed |
| Total cholesterol | LABA66 | mg/dL | 62 | 587 | Log-transformed |
| Triglycerides | LABA67 | mg/dL | 20 | 6366 | Log-transformed |
| HDL-cholesterol | LABA68 | mg/dL | 9 | 141 | Log-transformed |
| LDL-cholesterol | LABA69 | mg/dL | 22 | 417 | Log-transformed |
| Weight | ANTA4 | kg | 34.5 | 189.2 | Log-transformed |
| Waist girth | ANTA10A | cm | 31 | 202 | Log-transformed |
| Hip girth | ANTA10B | cm | 35 | 306 | Log-transformed |
| Average systolic | SBPA5 | mmHg | 74 | 232 | Log-transformed |
| Average diastolic | SBPA6 | mmHg | 39 | 135 | Log-transformed |
| Insulin post OGTT | INSULIN_OGTT | calibrated to mU/L | 1.5 | 960.5 | Log-transformed |
| White blood count | LABA1 | count $\times 10^9$ | 1.2 | 31.8 | None |
| % Neutrophils | LABA10 | percentage | 0 | 92 | None |
| % Lymphocytes | LABA11 | percentage | 5 | 90 | None |
| % Monocytes | LABA12 | percentage | 0 | 47 | None |
| % Eosinophils | LABA13 | percentage | 0 | 36 | None |
| % Basophils | LABA14 | percentage | 0 | 5.5 | None |
| Red blood cell count | LABA2 | count $\times 10^{12}$ | 2.21 | 7.61 | Log-transformed |
| Hemoglobin | LABA3 | g/dL | 4.3 | 20.6 | Log-transformed |
| Platelet count | LABA9 | count $\times 10^9$ | 26 | 874 | Log-transformed |
| Alanine Aminotransferase | LABA74 | U/L | 5 | 651 | Log-transformed |
| Aspartate Aminotransferase | LABA75 | U/L | 5 | 515 | Log-transformed |
| Gamma-Glutamyl Transferase (GGT) | LABA102 | U/L | 2 | 1660 | Log-transformed |
| Ferritin | LABA103 | ug/L | 7 | 12745 | Log-transformed |
| Iron | LABA82 | ug/dL | 8 | 320 | Log-transformed |
| Apnea/Hypopnea Index | SLPA54 | events/hour | 0 | 141.67 | Log-transformed |

## 3 Results

### 3.1 Theoretical cause of inflation in association meta-analysis due to between-study relatedness

For the following derivations, we will assume the fixed-effects meta-analysis framework implemented by METAL [28]. Suppose that there are *S* studies we wish to meta-analyze, each study *j* ∈ {1, …, *S*} with estimated coefficients 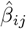 and standard errors 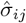 at SNP *i* The z-scores are

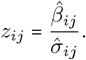

When the individual studies are calibrated, then under the null hypothesis of no association we have that E [*z*_*ij*_] = 0 and Var (*z*_*ij*_) = 1. The combined z-score *z*_*i*_ is the average per-study z-score weighted by the per-study standard error, namely

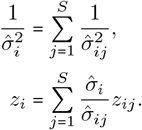

When studies are calibrated and independent of each other, the combined z-score is also calibrated. In particular, treating *z*_*ij*_ as random but 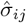 as fixed for simplicity, it follows under the null hypothesis that

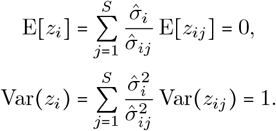

However, when studies have correlated statistics, the combined variance exceeds 1, making these statistics inflated even if the individual studies are calibrated:

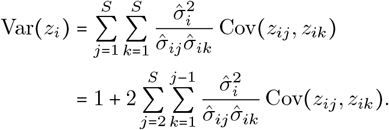

In Appendix A we derive a complete form for Cov (*z*_*ij*_, *z*_*ik*_) under a joint LMM as the true model, which has a complicated dependence on the kinship matrix between studies *j* and *k*, the trait heritability, and the model trait covariance matrices **V**_*j*_ of each separate study *j*. Treating the relatedness values between studies as random and independent, then the correlation term that leads to inflation has the approximate form

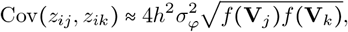

where 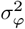 is the variance in between-study kinship values, and *f* (**V**_*j*_) ≥ *n*_*j*_ is a function (see Appendix A for closed form) that empirically scales with study *j* sample size (linearly for population structure, superlinearly under within-study cryptic relatedness), and further increases with heritability and study *j*’s *F*_ST_ (Figure 3). Each of these factors is non-negative, proving that relatedness between studies results in inflation upon meta-analysis. Further, for a fixed amount of cryptic relatedness variance between studies 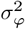, set indirectly via a threshold on kinship estimates for example, inflation will grow with sample size, so very distant relatives could result in substantial inflation for large, biobank-scale datasets.

**Fig. 3:**
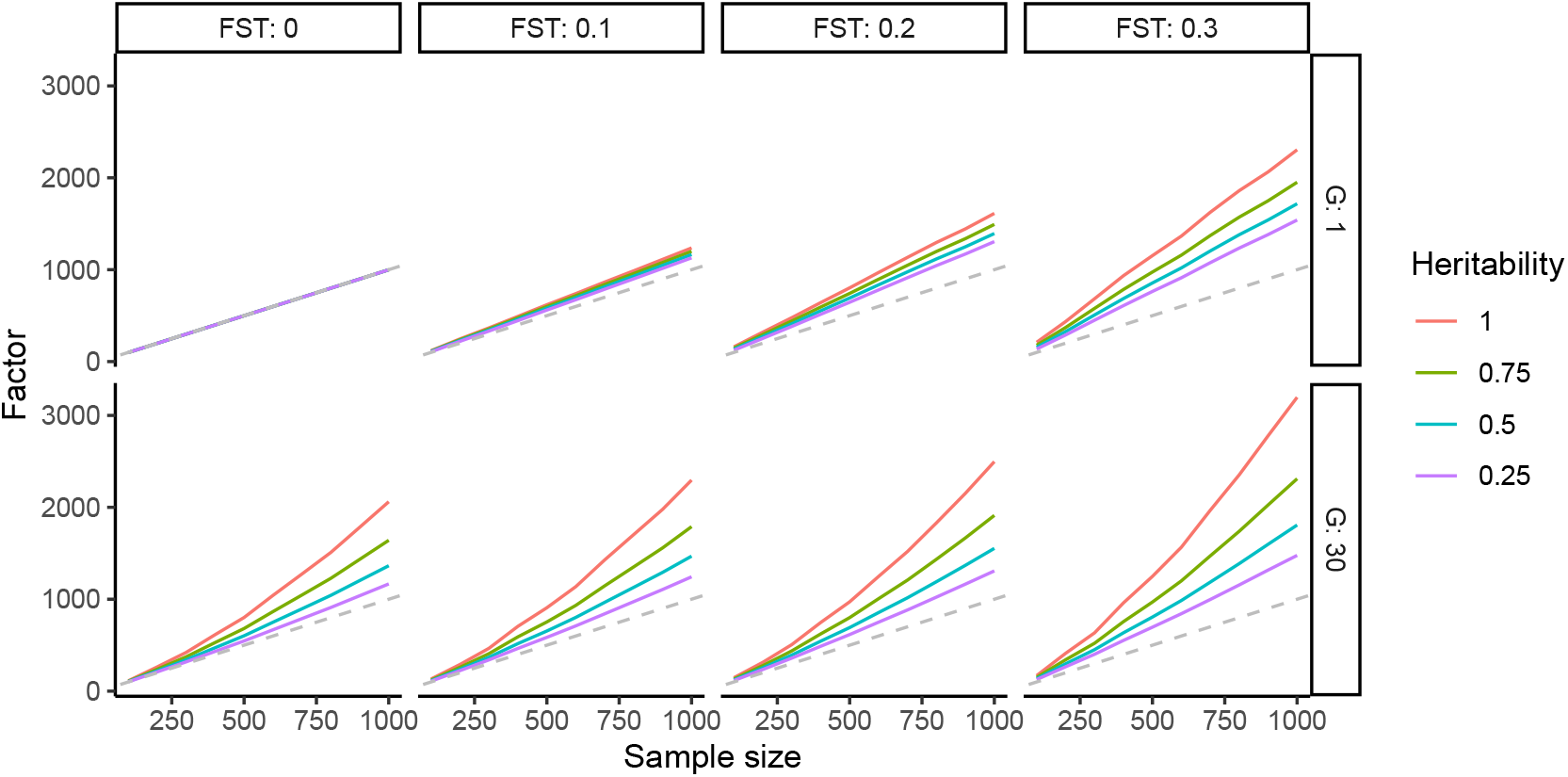
Dependence of within-study factor on sample size, population and family structure, and heritability. We plotted *n*_*j*_ versus the within-study factor *f* (**V**_*j*_) (see Appendix A) for a single study *j*, separately for combinations of *F*_ST_, heritability, and family structure (*G* = 1 versus 30 generations). Dashed gray line is *y* = *x*. For each combination of *G* and *F*_ST_, kinship matrices were constructed for *n* = 1, 000 individuals from the 1-dimensional admixture model used in [29] followed by *G* generations of family structure (**Methods**), then 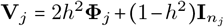 was calculated according to the desired heritability. Lower sample sizes were obtained by subsampling.

### 3.2 Meta-analysis inflation caused by between-study cryptic relatedness confirmed in simulated data

In all of our evaluations, we contrast three types of studies. The joint analysis fully accounts for population structure and cryptic relatedness and is taken as the baseline for good performance. The subpopulation-stratified meta-analysis (subpop-meta) is a common study design where individual studies are performed in single populations before combining resulting summary statistics for meta-analysis. Subpop-meta, also known as trans-ethnic meta-analysis, potentially handles cryptic relatedness successfully as distant family members are expected primarily within subpopulations rather than across them. Lastly, the sex-stratified meta-analysis (sex-meta) has the opposite property, namely that it is expected to result in considerable cryptic relatedness between studies (that is, between male and female subsets) to the degree that it is present in the study overall. Sex was chosen partly because it has been suggested in the literature [13], but also for convenience. In practice it is not important that the meta-analysis is sex-stratified, as sex is random compared to all other variables. Many other designs can result in similarly extreme cryptic relatedness between studies, particularly separate studies of the same population.

We ran 20 replicates of simulated genotypes and both binary and quantitative traits across the 4 scenarios (Figure 2) and evaluated performance for joint GWAS and sex and subpopulation stratified meta-analyses. Simulation 1 has population structure but no family structure, and is treated as the baseline against which the effects of cryptic relatedness are compared, since the rest contain family structure. In simulation 1, subpop-meta represents common trans-ethnic meta analyses, whereas sex-meta is a meta-analysis of individually multiethnic studies. Simulation 2 adds 30 generations of family structure only within populations. Subpop-meta is another common design for studies each of one of a few distant global populations, so cryptic relatedness is adequately modeled within studies, while no cryptic relatedness exists between studies. Simulation 3 simulates admixture across populations, but family structure remains relatively localized so it is expected to continue to have a larger effect for sex-meta versus subpop-meta. Lastly, simulation 4 is like simulation 3 without the population structure, for another comparison between the two effects. Simulation 4 has artificial subpopulations constructed by grouping individuals consistent with their family structure, which again yields minimal family structure between studies for subpop-meta compared sex-meta.

In simulations with family relatedness (simulations 2, 3, 4), we see both severe inflation (*λ* >1.05 and SRMSD_*p*_ >0.01) and reduced AUC_PR_ for sex-meta compared to subpop-meta and joint analysis (Figures 4 and 5, for binary and quantitative traits, respectively). When family relatedness exists across subpopulations (simulations 3, 4), we observe higher inflation in sex-meta compared to when family relatedness exists only within subpopulation (simulation 2).

**Fig. 4:**
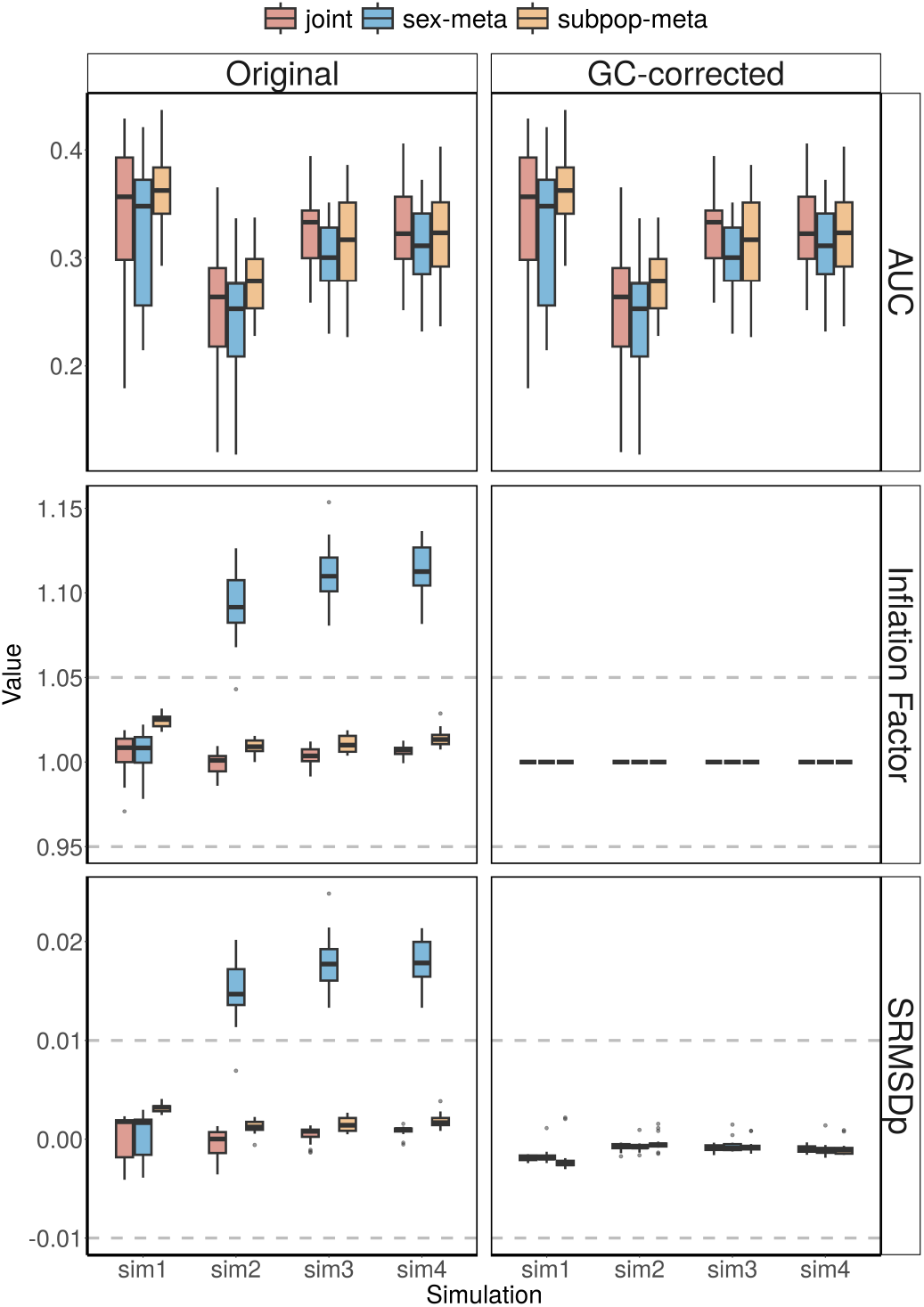
Simulation results confirm cryptic relatedness between studies results in considerable inflation upon meta-analysis. Simulations (x-axis) have different presence and arrangement of population and family structure. AUC_PR_ reflects calibrated power (higher is better), while both inflation factor (1 is best, >1.05 is inflated) and SRMSD_*p*_ (0 is best, >0.01 is inflated) measure null p-value calibration, the latter more strictly (see **Methods**). Results from 20 replicates using binary trait LMMs show that simulations with 30 generations of family relatedness (sim 2,3,4) demonstrate severe inflation and SRMSD_*p*_ values for sex-meta analyses and overall lower AUC_PR_ compared to subpop-meta and joint analyses. GC-corrected results show that despite improved type I error control (inflation factor and SRMSD_*p*_), the correction does not improve the loss of calibrated power (AUC_PR_) due to confounding.

**Fig. 5:**
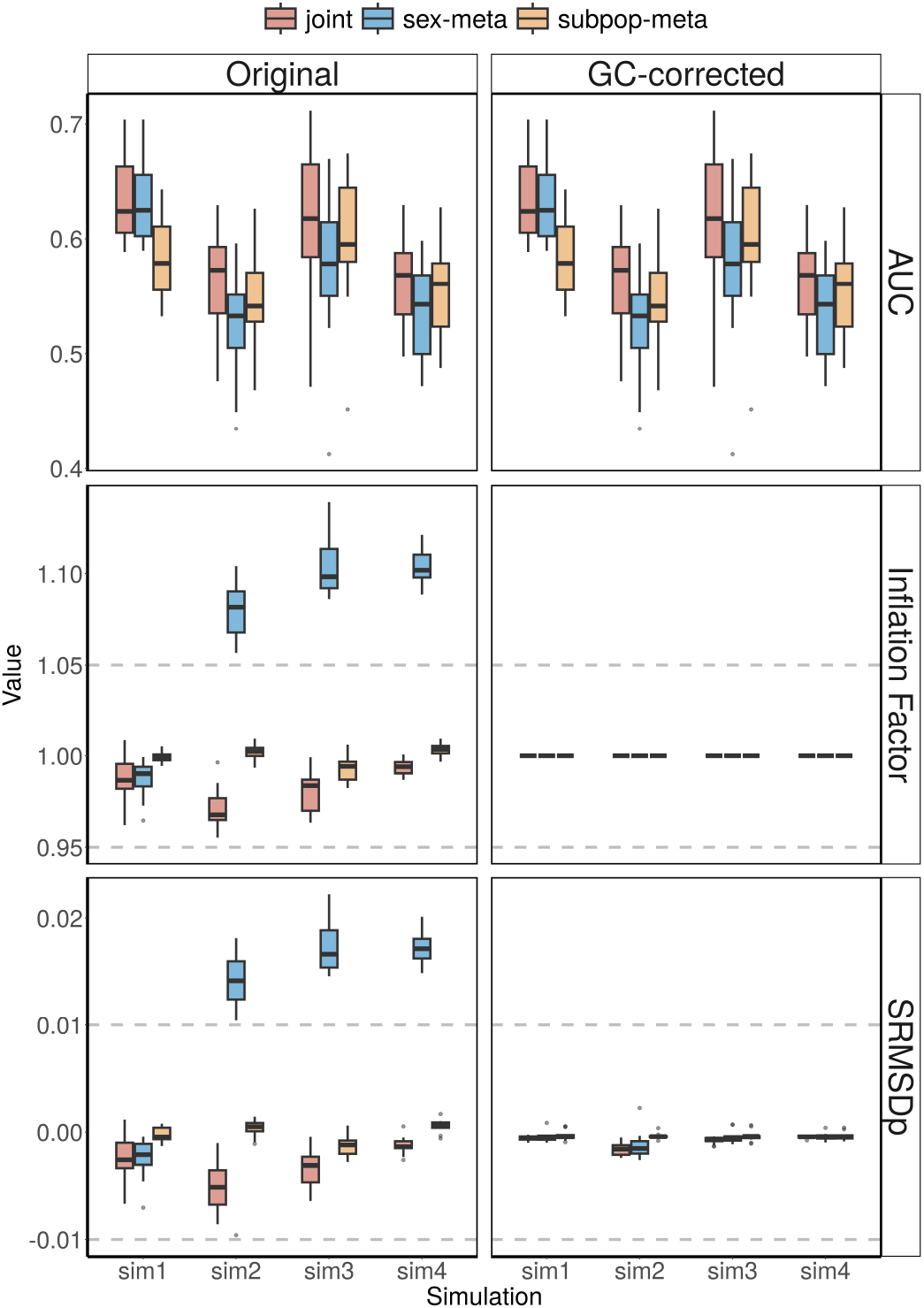
Simulation results for quantitative traits. Results from 20 replicates using a quantitative trait LMM exhibit similar patterns as the binary model (Figure 4) across all four simulation scenarios, where GC-corrected results improve inflation factor and SRMSD_*p*_, but not AUC_PR_.

Remarkably, unlike family structure, differences in ancestry (population structure) between studies appears to have a negligible effect on inflation and power upon meta-analysis, particularly for simulations 1-3 where it would have caused subpop-meta to differ more from joint analysis. Instead, joint is only slightly more calibrated than subpop-meta in binary traits, though the reverse holds in quantitative traits (both are considered effectively calibrated in all of these cases), and there is no clear difference in power between them except for one case: simulation 1 for quantitative traits only, where joint substantially outperforms subpop-meta. There is also no clear difference in performance between cases where subpopulations are entirely within studies (subpop-meta simulations 1-2) or when several subpopulations are present within each study so there is structure but not relatedness across studies (sex-meta for simulation 1), and when there is admixture so ancestries are mostly but not exclusively within single studies (subpop-meta simulation 3), compared to their joint counterparts or when there is no population structure (subpop-meta simulation 4). Next, we consider the GC correction, which by construction always results in perfect inflation factors of one (Figures 4 and 5). For that reason, here in particular we relied on the stricter SRMSD_*p*_ statistics to determine if GC actually results in improved null p-value calibration. Remarkably, GC correction consistently resulted in ∣SRMSD∣_*p*_ < 0.01, which is considered calibrated [12], for all approaches, including sex-meta that was highly inflated before the correction. Nevertheless, AUC_PR_ is unaffected by GC, because GC does not rerank SNPs while AUC_PR_ is a function of only SNP ranks and their true classes (causal or null). Overall, we find that in these scenarios GC successfully corrects the type 1 error rate control that otherwise results from inflation, but it does not correct the loss of calibrated power due to confounding by cryptic relatedness.

### 3.3 Inflation in meta-analyses of real data

We analyze two real datasets, with real genotypes and phenotypes, to determine if the effects and magnitudes observed in our simulations are also present in practice, as well as whether our GC solution works in practice. However, since true causal variants are generally unknown for real phenotypes, it is not possible to calculate AUC_PR_ and SRMSD_*p*_ in these cases, so we focus on inflation factors and quantile-quantile (QQ) plots.

First we considered T2D-GENES SAMAFS, a study that is large for a family study, but small compared to population studies (*n* = 914 individuals). Since there will be considerable family relatedness between the male and female subsets, this dataset produces a clear demonstration of the inflation in sex-stratified meta-analysis that we already illustrated with our simulations. We compare the inflation factors between the joint GWAS performed by SAIGE and the sex-stratified meta-analysis performed by METAL. All 21 traits exhibited severe inflation factors in the meta-analysis, with values exceeding the standard 1.05 threshold except for one case near the threshold, cystatin C, which had *λ* = 1.042 (Figure 6). In contrast, all joint analyses were calibrated (0.95< *λ*< 1.05), although two traits were near each of the thresholds: height (*λ* = 0.952) and cholesterol (*λ* = 1.047).

**Fig. 6:**
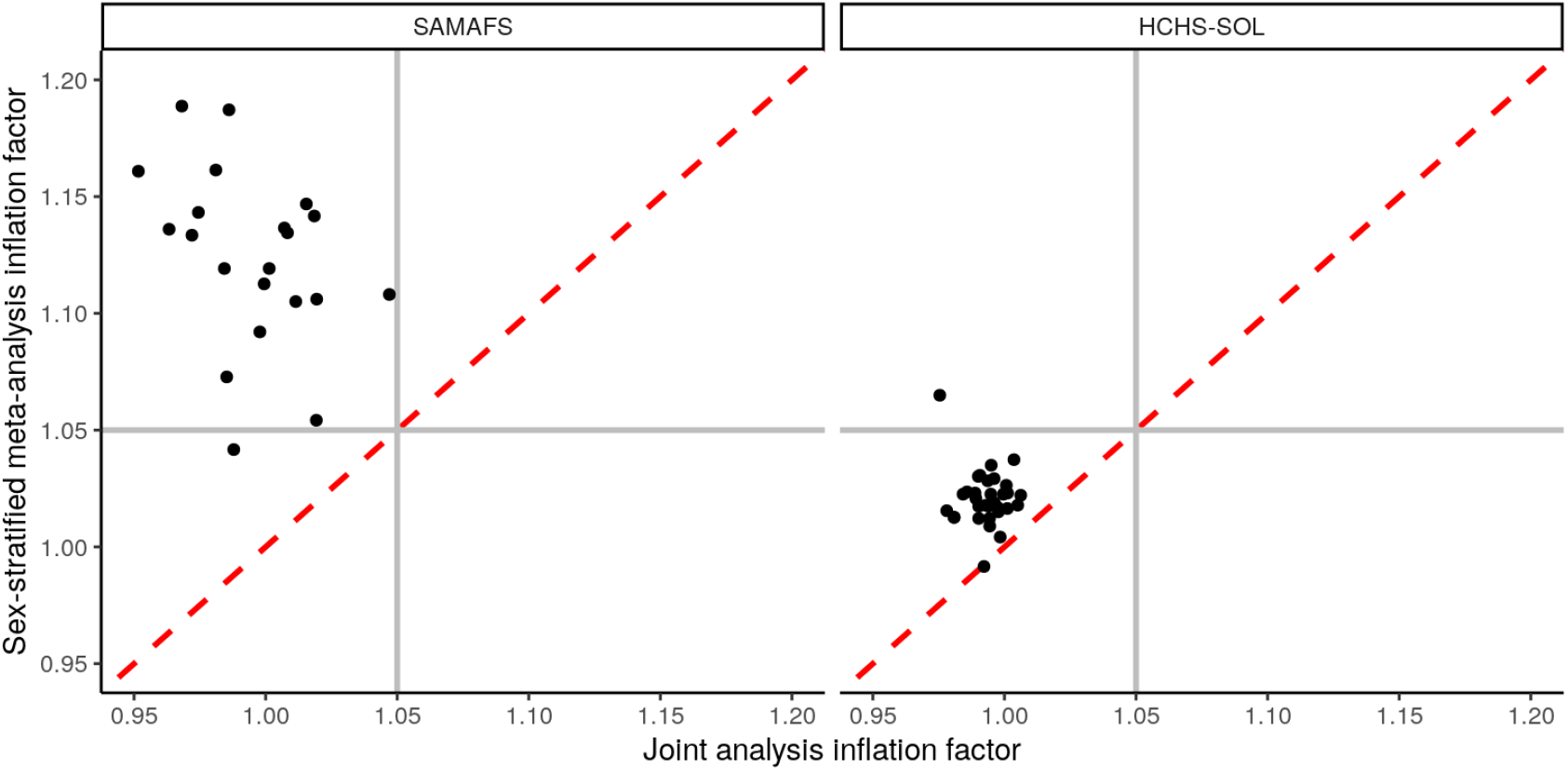
Inflation greater for sex-stratified meta-analyses of real genotypes and phenotypes. Inflation factors in joint analysis (x-axis) are smaller than those of sex-stratified meta-analysis (y-axis) for all but one of the traits analyzed. Values greater than 1.05 (horizontal and vertical gray lines) are considered inflated. Red dashed line is *y* = *x* line. In both datasets, all traits were calibrated in the joint analysis. In the T2D-GENES SAMAFS family study (20 quantitative traits and the single binary trait), all traits except one are inflated under the meta-analysis approach. In the HCHS/SOL population study (33 quantitative traits), only one trait is inflated under meta-analysis, although inflation factors were still consistently larger for meta-analysis compared to joint analysis.

Next we considered HCHS/SOL, which is a large population study of Hispanic individuals (*n* = 11,721 individuals), which did not specifically sample close individuals, but nevertheless likely contains cryptic relatedness as do most large studies. Only one of the 33 quantitative traits tested, height, exhibits a high inflation factor of 1.065 in the sex-stratified meta-analysis. Nevertheless, calibration is exceptional under the joint analysis, and all but one trait (iron blood concentration) show a higher inflation factor in sex-stratified meta-analysis compared to joint analysis (Figure 6). Thus, while the effect of cryptic relatedness is less severe in this population study, it is still measurable and highly consistent across traits.

Lastly, we consider the effect of GC correction of inflated meta-analysis p-values of the real phenotypes. As explained earlier, it is not possible to evaluate GC using inflation factors *λ*, since GC always results in *λ* = 1, while that does not guarantee adequate type I error control at other quantiles. Absent more precise measures such as SRMSD_*p*_, which is only available for simulated datasets with known causal variants, we performed a more holistic assessment by visualizing calibration across quantiles using QQ plots. For simplicity, we focused on the top three most inflated traits per dataset, which will have larger corrections performed by GC and thus a greater potential for overcorrection. We again found good performance for GC correction of meta-analyzed p-values, which are visually close to the null expectation across a large portion of the curves, especially on the lower end as desired (Figure 7). In contrast, inflation for the SAMAFS traits is visually noticeable, whereas HCHS/SOL has negligible inflation, as our calculated inflation factors determined earlier. If GC had overcorrected or otherwise failed, since GC forces the median to go through the diagonal (null expectation), we would expect at least a portion of the QQ plot curve to fall below the diagonal, especially among the lower -log10(p) values, but this did not occur. Therefore, as observed earlier for the simulated data, GC appears to be an adequate solution to correct inflation, but not loss of power, in meta-analysis due to cryptic relatedness.

**Fig. 7:**
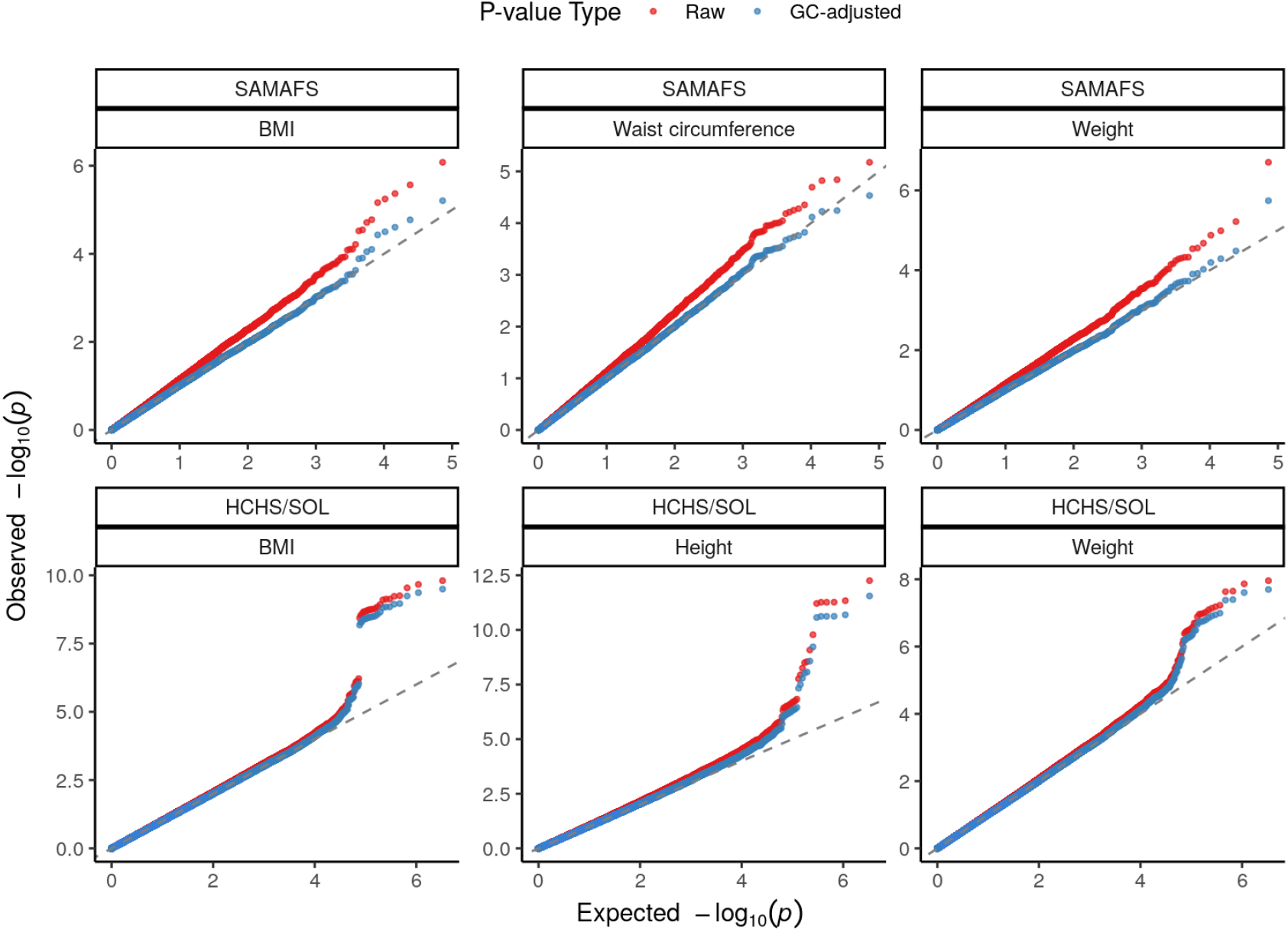
Quantile-quantile plots for meta-analysis p-values of most inflated traits before and after GC correction. Top three traits per dataset ranked by meta-analysis inflation factor. In SAMAFS, original (raw) inflation is severe (curve departs early from the null expectation, which is the diagonal *y* = *x* line, sits primarily above it), and GC successfully corrects inflation (curve follows diagonal much longer) without overcorrecting (curve does not dip below diagonal at the lowest values). In HCHS/SOL, GC also succeeds without overcorrecting, although original inflation was less severe.

## 4 Discussion

Meta-analysis is a well-established GWAS approach that aggregates data from multiple smaller studies to achieve a larger effective sample size and utilizes summary statistics instead of individual-level genotype data to improve power and accessibility. The main drawback is that the independence assumption between studies is not always met, and the effect of cryptic relatedness on GWAS meta-analysis was unclear before our work. Using simulations and real data, we confirmed that the presence of family structure between studies causes inflation upon meta-analysis, which is severe for family studies at low sample sizes, and observed with a smaller effect in population studies with up to *n* ≈ 10, 000 individuals. Both binary and quantitative traits GWAS models show strong inflation in sex-stratified meta analysis. Our theory predicts that, if there is cryptic relatedness between studies, increasing sample size also increases confounding (Figure 3), which intuitively is due to the higher number of related individuals in the sample population, for the same levels of relatedness. Even at low sampling fractions, it has been shown that a substantial fraction of sampled individuals are expected to have at least one second cousin within the sample [32]. Thus, we believe that accounting for cryptic relatedness between large cohort studies, such as biobanks, will be a crucial challenge that existing meta-analysis approaches are ill-equipped for.

Although cryptic relatedness between studies causes inflation in meta-analysis, population structure (ancestry differences) between studies does not appear to have the same effect in our simulations. Our hypothesis is that these cases behave differently due to the dimensionality of their structures. Population structure is low dimensional because it is widely shared, so modeling it within studies (via the intercept for single ancestries, or PCA for more complex admixture scenarios) suffices to completely remove the effect between studies, resulting in test statistics that are independent between studies. On the other hand, cryptic (family) relatedness is high dimensional, and it is not broadly shared. In that case, since the relationship between a pair of individuals from different studies is by hypothesis absent within studies, it is not modeled by them and remains a confounder upon meta-analysis. Thus, our results suggest that meta-analyses of studies from different populations, especially distant populations with less migration between them, are likely much less affected by cryptic relatedness and thus less likely to experience inflation, even in the presence of ancestry differences, absent other causes.

Recent work that suggested applying sex-stratified meta-analysis to correct for participation bias did not consider relatedness between individuals of opposite sex drawn from the same population [13]. Although we suggested that a joint analysis is preferable to a sex-stratified meta-analysis to avoid inflation due to cryptic relatedness, we have not addressed the primary concern of that work, which is that there is a potentially sex-specific effect in the observed association, which is due to sex-differential participation bias that we wish to ignore [33]. Meta-analysis in this case allows each sex to have its own intercept as well as its own genetic effect coefficient, so it successfully makes sex-specific associations vanish. A standard joint analysis allows for a sex-specific intercept by including sex as a fixed covariate, but this does not allow for sex-specific genetic effects unless we introduce an interaction term between genotype and sex, and we condition the overall association (the genetic effect shared between men and women) by the sex-specific effects. The most popular scalable LMM packages do not allow for such interactions between genotypes and covariates, but it is possible to carry out such analyses in custom LMM implementations, although it would be preferable for highly optimized versions to be available.

There are three potential solutions to the problem of between-study cryptic relatedness: (1) pruning of cryptic relatives between studies, (2) correcting the resulting inflation due to meta-analysis, or (3) applying joint analysis with a method that models cryptic relatedness such as LMMs. For the first option, methods like KING, GERMLINE, TRUFFLE, or IBIS [34–37] can infer the degrees of relatedness for filtering as a quality control step. However, removing increasingly distant relatives can result in large sample size reductions: In datasets such as Human Origins, HGDP, and 1000 Genomes, removing 4th degree relatives reduces sample sizes by 5% to 10%, yet confounding due to cryptic relatedness remained for non-LMM methods, showing that the larger number of very distant relatives pose significant problems when they are not modeled in GWAS [12]. For the second option, Genomic Control (GC) allows for correction of inflated test statistics, which in both our simulated and real data results in remarkably accurate type I error control. However, GC does not improve the ability to detect true signals compared to the joint analysis (Figures 4 and 5), which makes sense since GC does not directly model cryptic relatedness. Consequently, we recommend avoiding meta-analysis of studies that are drawn from the same genetic populations, although when there is no other choice, GC may allow for adequate type I error control in those cases. For the third option, besides traditional joint LMM analyses, federated genome-wide association studies are another alternative to traditional meta-analysis that utilizes full genomic datasets instead of summary statistics for conducting collaborative GWAS across different institutes without jeopardizing patient privacy. Our work may motivate further developments of such federated approaches, or to improve responsible data sharing policies to facilitate joint analyses.

## Acknowledgments

This work was funded in part by the Duke University School of Medicine Whitehead Scholars Program, a gift from the Whitehead Charitable Foundation.

We would like to thank the staff and participants of the SAMAFS and HCHS/SOL study for their important contributions. The research reported in this article was supported by National Institutes of Health grants. The genetic and phenotypic data were provided by the San Antonio Family Heart Study (SAFHS) investigators and supported by the National Heart, Lung, and Blood Institute (NHLBI) [P01 HL045222] and the San Antonio Family Diabetes/Gallbladder Study (SAFDGS) investigators and supported by the National Institute of Diabetes and Digestive and Kidney Diseases (NIDDK) [R01 DK047482, R01 DK053889]. The phenotypic data were also provided by the Veterans Administration Genetic Epidemiology Study (VAGES) investigators and supported by the Health Services Research and Development, U.S., Department of Veteran Affairs and the Family Investigation of Nephropathy and Diabetes (FIND) - San Antonio (FIND-SA) Component and its extension called the Extended FIND (E-FIND) [U01 DK57295] investigators and supported by the NIDDK. The SAFHS gene expression assays were supported by a donation from the Azar and Shepperd families. The exome sequencing, exome chip genotypic, and whole genome sequencing data were provided by the T2D-GENES Consortium grants U01 DK085524, U01 DK085584, U01 DK085501, U01 DK085526, and U01 DK085545 and investigators supported by the NIDDK. This manuscript was not prepared in collaboration with investigators of the T2D-GENES SAMAFS/Consortium and does not necessarily reflect the opinions or views of the members of the T2D-GENES SAMAFS/Consortium, or the NIDDK. The Hispanic Community Health Study/Study of Latinos is supported by contracts from the National Heart, Lung, and Blood Institute (NHLBI) to the University of North Carolina at Chapel Hill, Chapel Hill, NC (N01-HC65233), University of Miami, Miami, FL (N01-HC65234), Albert Einstein College of Medicine, Bronx NY (N01-HC65235), University of Illinois, Chicago IL (N01-HC65236), and San Diego State University, San Diego CA (N01-HC65237). The following Institutes/Centers/Offices contribute to the HCHS/SOL through a transfer of funds to the NHLBI: National Center on Minority Health and Health Disparities, the National Institute of Deafness and Other Communications Disorders, the National Institute of Dental and Craniofacial Research, the National Institute of Diabetes and Digestive and Kidney Diseases, The National Institute of Neurological Disorders and Stroke, and the Office of Dietary Supplements.

## Data and code availability

The data and code generated during this study are available on GitHub at https://github.com/OchoaLab/meta-gwas-cryptic-inflation

## Appendices

### A The covariance between the z-scores of two GWAS with relatedness between them

#### A.1 General formula under joint LMM

Here we derive the form of Cov (*z*_*ij*_, *z*_*ik*_) under the assumptions of the joint LMM of studies *j* and *k*, which clearly reveals how relatedness between studies results in within-study z-scores that are correlated across studies. Specifically, we assume the following trait covariance matrix for studies *j* and *k* jointly, which is implied by Eq. (1), written to clarify its block structure:

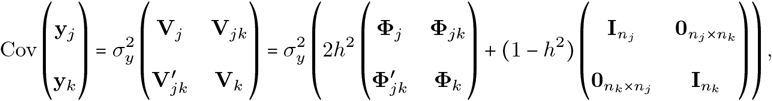

where **V**_*jk*_ and **Φ**_*jk*_ are *n*_*j*_ × *n*_*k*_ trait covariance and kinship matrices between the individuals in studies *j* and *k*, respectively, 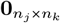 is an *n*_*j*_ × *n*_*k*_ matrix of zero values, and the rest follows the notation in the main methods. Thus, subsetting to the between-studies block, we obtain the cross-covariance between the trait vectors in two different studies:

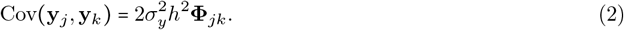

The joint LMM assumes that the trait is fundamentally the same across studies, so 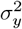 and *h*^2^ are the same too. Furthermore, we assume there is no additional covariance due to non-genetic environment that is unmodeled; however if that were present, it could increase inflation further, so that simplistic assumption does not negate our key conclusion.

Given a known **V**_*j*_, the standard generalized least squares solution used by modern LMMs for the genetic effect and its estimated variance for SNP *i* and study *j* are

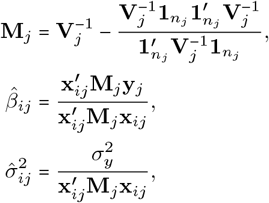

where for notational simplicity we assumed that no covariates beyond the intercept were present, though the key results hold without this restriction. Note **M**_*j*_ is symmetric since **V**_*j*_ is too. The z-score is therefore

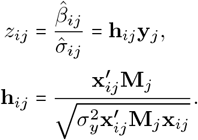

Note **h**_*ij*_ does not depend on the trait so it is considered fixed for now. Therefore, the covariance between test statistics across studies *j* and *k* arises from the covariance between their traits, which is in turn given by the relatedness between studies as stated in Eq. (2):

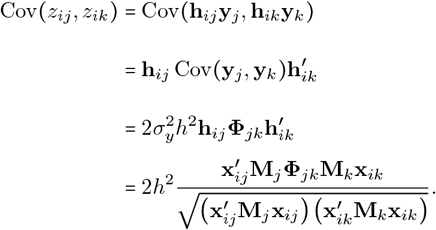

Furthermore, 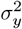 no longer appears in these formulas, as inflation does not depend on the scale of the trait, but it does depend on the heritability of the trait.

#### A.2 ormula marginalizing genotypes under kinship model

In order to understand the connection between allele frequency, population structure, and inflation, it is further necessary to take expectations of the above covariance over the possible values of the genotype vectors. For this, we consider the kinship model, which is also the motivation behind the LMM [7]. Stated to cover the genotypes of both studies *j* and *k*, the kinship model establishes that the mean and covariance structure of these genotypes is given by

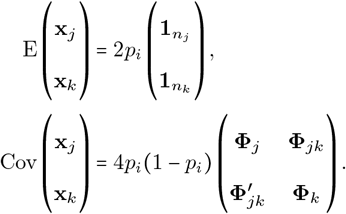

Using the well-known identity

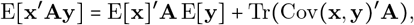

where **A** is fixed, we calculate the expectation over random genotypes of the numerator and the parts of the denominator of the desired covariance:

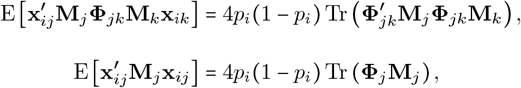

where the mean terms vanish since the intercept 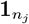 is in the null space of **M**_*j*_ and the same for study *k*:

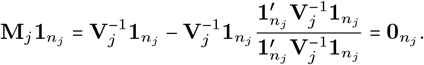

Thus, assuming we can approximate the expectation of the function of those random variables with the function of their expectations, we get an approximate expectation over genotypes of

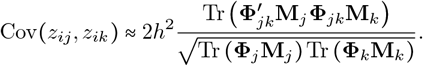

Note that this inflation term therefore does not depend on the allele frequency *p*_*i*_ of the tested locus, as those factors have canceled out.

#### A.3 Formula under random kinship distribution

The previous formulas for specific kinship matrices can be easier to understand if we treat kinship values themselves as random. To calculate an expectation, we simply need mean and variance of the distribution of cross-kinship values, and for simplicity we will assume that different values ***Φ*** in the cross-kinship matrix are uncorrelated of each other. First note this identity if the elements of the *m* × *n* matrix **X** are independent and identically distributed with variance *σ*^2^, while **A** is fixed, then

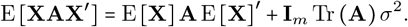

Thus, applied to our case, which has 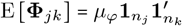 and variance 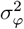, which is appropriate for cryptic relatedness but not population structure, we get

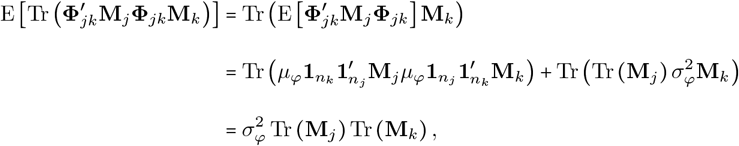

where the mean terms vanish again as before.

The kinship within studies is more challenging to model as random because kinship is also estimated for the LMM so it appears in the **M**_*j*_ terms as well (in contrast, kinship between studies does not appear in within-study LMM parameters). Nevertheless, since 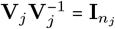, and expanding the uninverted **V**_*j*_, we get a way to rewrite one product involving **Φ**_*j*_ into terms that do not involve this kinship except through 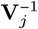:

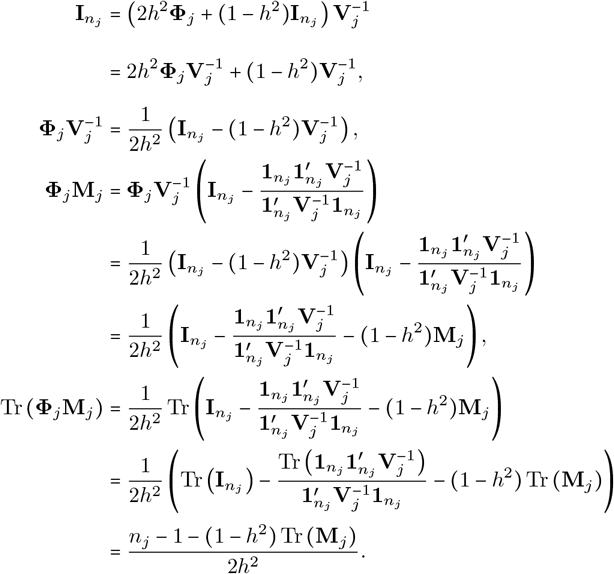

The above assumes *h*^2^ ≠ 0, which is also necessary to have inflation. All together, the z-score covariance becomes

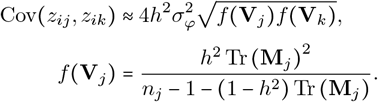

